# Role of the right middle occipital gyrus in egocentric spatial orientation in reference to gravitational information: Evidence from a pre-registered rTMS study

**DOI:** 10.1101/2024.03.07.584011

**Authors:** Keisuke Tani, Eiichi Naito, Koji Mizobe, Satoshi Hirose

## Abstract

Accurate perception of the orientation of external objects relative to the body, known as *egocentric spatial orientation*, is fundamental to action. Previously, we found via behavioral and magnetic resonance imaging voxel-based morphometry studies that egocentric spatial orientation is distorted when the whole body is tilted with respect to gravity, and that the magnitude of this perceptual distortion is correlated with grey matter volume in the right middle occipital gyrus (rMOG). In the present pre-registered study, we demonstrated that neural processing in the rMOG is indeed a cause of the perceptual distortion. We transiently suppressed neural activity in the rMOG by applying low-frequency repetitive transcranial magnetic stimulation (rTMS) and evaluated the consequent effect on perceptual distortion. Our results showed that while rTMS over the rMOG significantly reduced perceptual distortion when the body was tilted with respect to gravity, it did not affect egocentric spatial orientation when in the upright position. No changes in perceptual distortion were observed when rTMS was applied to a control site (right temporoparietal junction) or to air (sham TMS). These results indicate that neural processing in the rMOG during body tilt is an essential cause of perceptual distortion, suggesting that the rMOG is engaged in egocentric spatial orientation concerning gravitational information.

**Significance statement:** The findings of this pre-registered study support a causal role of neural activity in the right middle occipital gyrus (rMOG) in the perceptual distortion of egocentric spatial orientation induced by whole-body tilt relative to gravity. We suppressed neural activity in the rMOG using low-frequency repetitive transcranial magnetic stimulation (rTMS) and measured perceptual distortion. We observed a significant reduction in perceptual distortion after rTMS over the rMOG, but not after control or sham rTMS. These results provide, for the first time, direct evidence of the engagement of the rMOG in egocentric spatial orientation in reference to gravitational information.

## Introduction

Accurate perception of the orientation of external objects relative to the body (*egocentric spatial orientation*) is critical to the planning and execution of goal-directed actions. Indeed, distorted perception of a target’s orientation with respect to body-centered coordinates can deteriorate the accuracy of body movements such as arm reaching (Tani et al., 2018) and postural control (Barra et al., 2009; Jamal et al., 2018).

The subjective visual body axis (SVBA) task, also referred to as the longitudinal body axis or Z-axis task, was developed to quantify egocentric spatial orientation (Barra et al., 2008, 2009; Clement et al., 2007, 2009; Ceyte et al., 2009, Mars et al., 2005; Tani et al., 2023b). In the SVBA task, participants are asked to align the direction of a visual line parallel to the longitudinal axis of their own body. Although this task does not require participants to refer to the direction of gravity, SVBA task performance is strongly affected by gravitational information. Specifically, the indicated direction of the visual line is biased towards the side to which the body is laterally tilted (tilt-dependent error, TE; Barra et al., 2008; Ceyte et al., 2007, 2009; McFarland and Clarkson, 1966; Tamura et al., 2017; Tani et al., 2018, 2023a, 2023b; Tani and Tanaka, 2021; Bauermeister, 1964). In a recent study (Tani et al., 2023b), we found that the TE was correlated with the perceived degree of body tilt relative to gravity, independent of the actual body tilt angle. This suggests that the brain refers to the perceived direction of gravity to compute the egocentric orientation of visual objects (Tani et al., 2023b, 2023c; also see Tarnutzer et al., 2012).

Recently, using voxel-based morphometry (VBM), we found that the grey matter volume in the right middle occipital gyrus (rMOG) and the TE were significantly correlated across individuals (Tani and Tanaka, 2021), indicating a potential role of the rMOG in egocentric spatial orientation with respect to the direction of gravity. Here, to build upon the previous study, we validated the causal relationship between neural processing in the rMOG and egocentric spatial orientation. We performed a non-invasive neuromodulation experiment using low-frequency (1 Hz) repetitive transcranial magnetic stimulation (rTMS). Given that low-frequency rTMS can induce transient suppression of regional neural activity (Romero et al., 2002; Fitzgerald et al., 2006), we hypothesized that rTMS over the rMOG would alter the TE in the SVBA task. Additionally, we assessed the effect of rTMS over the right middle temporoparietal junction (rTPJ) on SVBA task performance. Although the rTPJ is known to contribute to multisensory integration processing regarding spatial orientation (Fiori et al., 2015; Khearmand et al., 2015; Santos-Pontelli et al., 2015) and body representation (Tsakiris et al., 2008; Donaldson et al., 2015 for a review), our VBM study (Tani and Tanaka, 2021) revealed no association between rTPJ and TE in the SVBA task. Therefore, we expected that rTMS over the rTPJ would not affect the TE.

## Methods and Materials

### Participants

Twenty healthy right-handed participants (6 women) with a mean (± standard errors; SE) age of 20.3 (± 0.28) years were recruited via posters and the internet. No participants had a history of psychiatric or neurological disorders, and all were naïve with respect to the rTMS experiment. The number of participants was determined according to a previous rTMS study on spatial orientation by Fiori et al. (2015). Participant handedness was assessed using the FLANDERS handedness questionnaire (Nicholls et al., 2013). Prior to the experiment, written informed consent was provided by all participants in accordance with the Declaration of Helsinki (2013). The experimental protocol was approved by the ethics committee of Otemon Gakuin University. To enhance the transparency and reproducibility of the study (Nosek et al., 2018), the number of participants, protocol, and statistical analysis were pre-registered in the Open Scientific Framework (https://doi.org/10.17605/OSF.IO/FEU8T).

### Experimental procedure and task

Each participant completed 3 experimental sessions in which they were exposed to the rMOG, rTPJ, or sham rTMS condition (see below). The sessions were on different days that were separated by intervals of at least 5 days. On each experimental day, the participants performed the SVBA task before (Pre-rTMS phase) and after rTMS exposure (Post-rTMS phase). The order of the 3 rTMS conditions was counterbalanced across participants.

During the SVBA task sessions in the Pre- and Post-TMS phase, the participants sat on a tilting chair (SP-PS100-Z, Pair Support, Japan) that could be rotated in the frontal plane with a peak velocity of 4.48°/s and acceleration of 2.61°/s^2^. The participants wore a head-mounted display (HMD; Oculus Rift S, Meta, California, USA) device and held a numeric keypad (TK-TCM011SV, Elecom, Osaka, Japan). Their right and left thumbs were placed on specific keys so that they could press the keys without using visual cues. The HMD device (along with their head), trunk, and legs, were firmly secured to the seat with bands and a seat belt.

Figure 1A shows the procedure on each experimental day. The SVBA task session consisted of 9 blocks, including 3 upright (UR), 3 right-side-down (RSD), and 3 left-side-down (LSD) blocks. The block order was randomized. At the beginning of the RSD or LSD block, the chair was tilted 10 degrees to the right or left, respectively (“Body tilt” in Fig. 1A bottom). In the UR block, the chair was not tilted. The participants completed 10 trials of the SVBA task in each body position.

**Figure 1.**
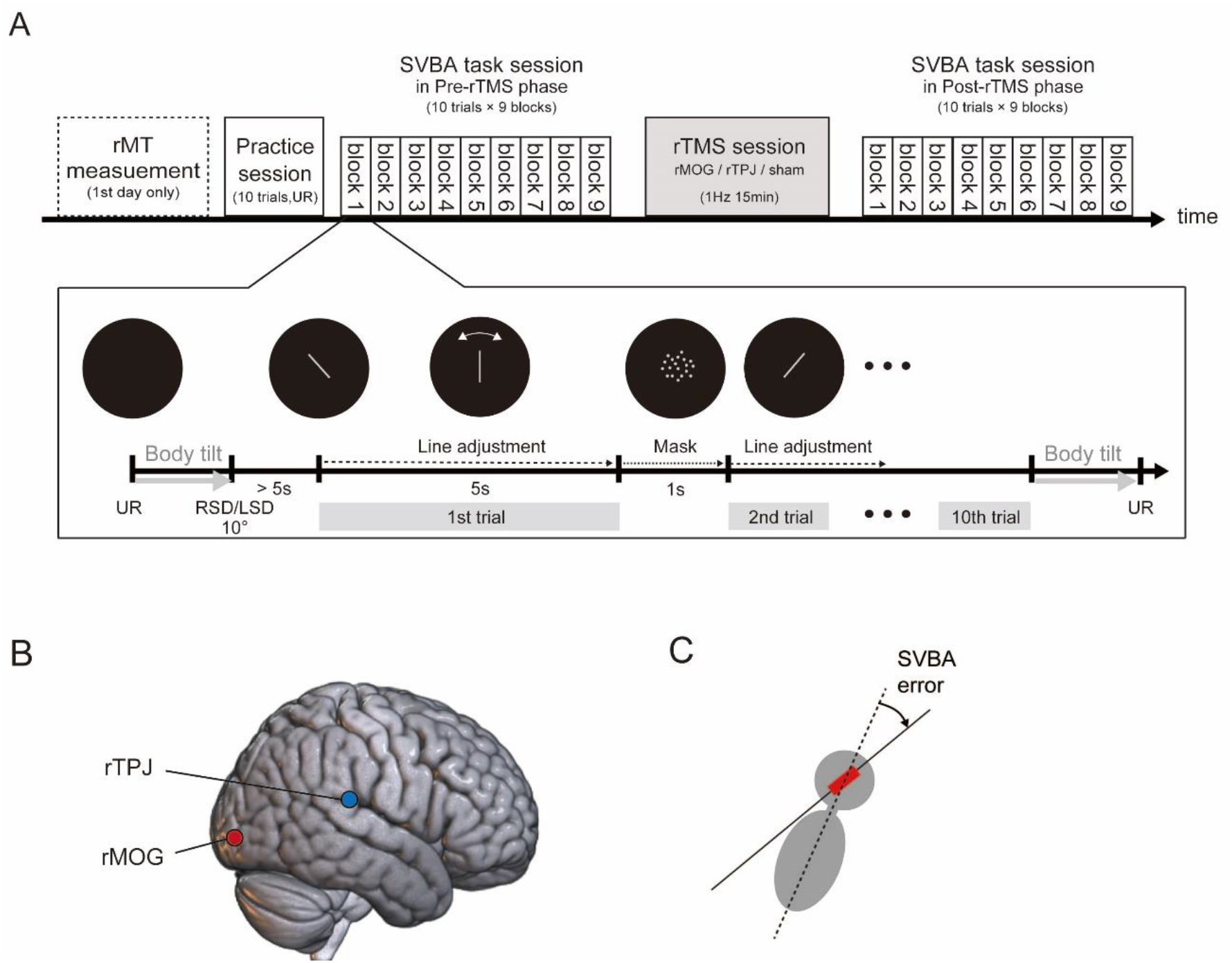
(A) Schematic overview of the experimental procedure on each day. In the SVBA task sessions in the Pre/Post-rTMS phase, the participants had 5s to align the orientation of the presented visual line so that it was parallel to the body longitudinal axis (“Line adjustment”). They performed 10 SVBA task trials in each block. A visual mask (random dots) was presented for 1s between each SVBA trial (“Mask”). The order of the body positions in the SVBA task was randomized. One of three TMS conditions was assigned for each SVBA task session, and the condition order was counterbalanced across participants. (B) The stimulation sites for the two active rTMS conditions. rTPJ, right temporo-parietal junction; rMOG, right middle occipital gyrus. (C) SVBA error calculation in each trial. We calculated the angular error between the actual body longitudinal axis (dotted line) and the indicated line (red bar and black solid line).

At the beginning of each trial, a grey line segment with a length corresponding to a visual angle of 53.1° was presented at the center of the display. The initial orientation of the line was randomly chosen to be ± 30°, ± 45°, or 90° relative to the vertical axis of the HMD. The participants were asked to use the numeric keypad to adjust this line parallel to their body’s longitudinal axis, and to make a response within 5s (“Line adjustment”). The inter-trial interval was set at 1s. During this interval, randomly positioned grey dots were presented within the circular area that would contain the bar (i.e. a circle with a diameter corresponding to a visual angle of 53.1°) to avoid low-level visual adaptation in the next trial (“Mask”; Clifford et al. 2007). To avoid semicircular stimulation caused by tilting the body, the first trial began more than 5s after the completion of the body tilt, as in a previous study (Tarnutzer et al., 2012). When the participants could not align the line within the time limit (5s) in a trial, they verbally reported their response to the experimenter immediately after the trial. After each block was complete, the chair orientation was returned to the upright position (“Body tilt”). To minimize the operational error and stabilize the data, each participant completed 10 trials of the SVBA task in the UR position (“Practice session” in Fig. 1A top) prior to the Pre-rTMS phase on each experimental day.

### rTMS

In the rTMS session, low-frequency (1 Hz) rTMS was applied to the participants for 15 min while they were seated and relaxed in a reclining chair. According to the rTMS condition on each experimental day, TMS was applied to the rMOG, rTPJ, or no brain region (sham TMS). TMS was delivered using a figure-eight coil with a wing diameter of 70 cm (MCF-B70; MagVenture, United Kingdom) connected to MagPro R20 magnetic stimulator (MagVenture, United Kingdom).

The coil position and orientation were manually adjusted using the Brainsight neuro-navigation system (Rogue Research Inc., Canada; Bashir et al. 2001) and anatomical T1-weighted brain images (MP-RAGE; repetition time [TR]: 1900 ms; echo time [TE]: 2.48 ms; inversion time [TI]: 900 ms; flip angle: 9°; matrix: 256 × 256; pixel size: 1.0 mm × 1.0 mm; slice thickness: 1.0 mm without inter-slice gap; number of slices: 208), which were obtained using a 3.0-T SIEMENS scanner (Vida, Germany) with a 64-channel head/neck coil. For the rMOG, we targeted the cortical surface using the following MNI coordinates: x = 35, y = −86, z = 6, according to our previous study (Tani and Tanaka, 2021). For the rTPJ, the target coordinates was x = 61, y = −37, z = 22 (Fiori et al., 2015; see Fig. 1B). The coil was placed tangential to the surface of the scalp and oriented with the handle pointing backward for the rMOG condition (Kassuba et al., 2014) and 45 ° upward for the rTPJ condition (Fiori et al., 2015). In the sham condition, the coil was placed in a cortical location identical to that in the rMOG condition, but held perpendicular to the scalp so that the brain was not stimulated.

The stimulus intensity of the rTMS was set at 90% of the individual resting motor threshold (rMT). The rMT was assessed on the first experimental day prior to the experiment (“rMT measurement” in Fig. 1A top). First, the motor-evoked potential (MEP) was recorded from the first dorsal interosseous (FDI) muscle of the left hand via Ag/AgCl surface electrodes. Electromyography (EMG) signals were amplified, bandpass filtered between 16–470 Hz, and sampled at 3 kHz with a Rogue EMG device. Then, the motor hotspot, the cortical spot where a TMS pulse evoked a MEP in the left FDI muscle with a maximum amplitude, was detected by applying single-pulse TMS to different positions around the right primary motor cortex. The rMT was defined as the minimum TMS current intensity that produced MEPs > 50 µV for at least 5 of 10 stimulation pulses (single-pulse TMS) applied to the motor hotspot during rest. The coil for the rMT determination procedure was oriented with the handle pointing backwards and laterally at a 45◦ angle away from the nasion-inion line (Mills et al., 1992). The mean (± SE) rTMS stimulus intensity was 50.4 (± 2.1) % of the maximum stimulator output.

### Data analysis

The present study had a repeated measures design with 3 within-subject factors: “body position” with 3 levels (UR, LSD, RSD), “rTMS condition” with 3 levels (rMOG, rTPJ, sham), and “phases” with 2 levels (Pre-, Post-rTMS). The measurement variable was the signed angular deviation of the indicated line from the actual body longitudinal axis in the SVBA task (SVBA error; Fig. 1C). Positive and negative values indicated the deviation towards rightward and leftward, respectively.

For each participant, we performed the following analysis. First, we excluded trials in which the participant could not align the line with the target orientation within the time limit (5s). Then, we averaged the SVBA error for the 30 trials in each condition. The effect of leftward or rightward body tilt on SVBA error (TE to the left and right: *TE*_L_, *TE*_*R*_, respectively) was quantified by subtracting the SVBA error in the UR position (denoted by *SVBAU*_*R*_) from that in the LSD or RSD position (*SVBA*_L*SD*_, *SVBA*_*RSD*_) as follows:

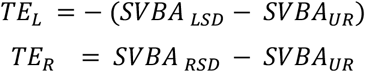

Note that the sign of *TE*_L_ was set opposite to that of *SVBA* _L_ − *SVBAU*_*R*_ so that a positive *TE*_L_ indicated that the shift of the indicated line was in the same direction as the body tilt, as well as *TE*_*R*_. The *TE*_L_ and *TE*_*R*_ were then averaged to obtain the *TE*. The *TE*_L_, *TE*_*R*_, and *TE* were calculated for each combination of phase and rTMS condition. Finally, for each rTMS condition, we calculated *ΔTE* by subtracting *TE* for the Pre-rTMS phase from that for the Post-rTMS phase. We considered *ΔTE* to be an indicator of the effect of rTMS on the distortion of egocentric spatial orientation induced by body tilt.

We first checked whether the SVBA error was biased in the direction of body tilt in the LSD and RSD positions, as was to be expected according to previous studies (Barra et al., 2008; Ceyte et al., 2007, 2009; McFarland and Clarkson, 1966; Tamura et al., 2017; Tani et al., 2018, 2023a, 2023b; Tani and Tanaka, 2021; Bauermeister, 1964).

After calculating *ΔTE* for each rTMS condition and each participant, we evaluated the group effects as follows. We performed a one-way repeated measures analysis of variance (ANOVA) of *ΔTE* after confirming the normality of the *ΔTE* distribution within each rTMS condition using the Shapiro-Wisk test (all *p* > 0.05). This was followed by paired Dunnett post-hoc tests (two-tailed) to compare *ΔTE* in the rMOG or rTPJ condition with that in the sham condition, regardless of the main effects revealed by the ANOVA, based on a previous statistics paper (Hothorn, 2016).

We then conducted two additional statistical analyses. First, we confirmed that the *TE* in the Pre-rTMS phase was not significantly different across the TMS conditions using a one-way repeated measures ANOVA and post-hoc tests (Dunnett). Second, we assessed the effect of rTMS on SVBA error in the UR position according to Δ*SVBAU*_*R*_, which was obtained by subtracting *SVBAU*_*R*_in the Pre-rTMS phase from that in the Post-rTMS phase. We calculated *ΔTE* and *ΔSVBAU*_*R*_ for each rTMS condition and each participant, and conducted a one-way (rTMS condition) repeated measures ANOVA followed by Dunnett post-hoc tests (rMOG/rTPJ vs. sham). The significance level was set at *p* < 0.05. All statistical analyses were performed using SAS statistical software (version 9.1; SAS, Inc., Cary, NC).

## Results

All participants completed all 3 experiments without any discomfort or adverse effects (e.g., seizure) during or after the stimulation trials. One participant (subject ID: 18) was excluded from the analysis because of a large trial-by-trial variability (standard deviation) in the *SVBAU*_*R*_ data that exceeded 3 SD from the mean of all participants. As a result, data from 19 participants were included in the group analysis. After excluding the trials in which the participants did not complete the task within the time limit (5s), data from 98.4% of the total trials were included in the analysis.

Figure 2A shows the mean SVBA error at each body position for each phase and rTMS condition. The SVBA error was nearly zero (overall mean = −0.80°) when the body was upright (UR). In contrast, as expected according to previous studies (Barra et al., 2008; Ceyte et al., 2007, 2009; McFarland and Clarkson, 1966; Tamura et al., 2017; Tani et al., 2018, 2023a, 2023b; Tani and Tanaka, 2021; Bauermeister, 1964), the SVBA was strongly biased in the direction of body tilt when the body was laterally tilted (LSD and RSD), regardless of the phase and rTMS condition. The consistency of this trend across individuals was confirmed by the positive *TE* values in all conditions in almost all participants (see Fig. 2B).

**Figure 2.**
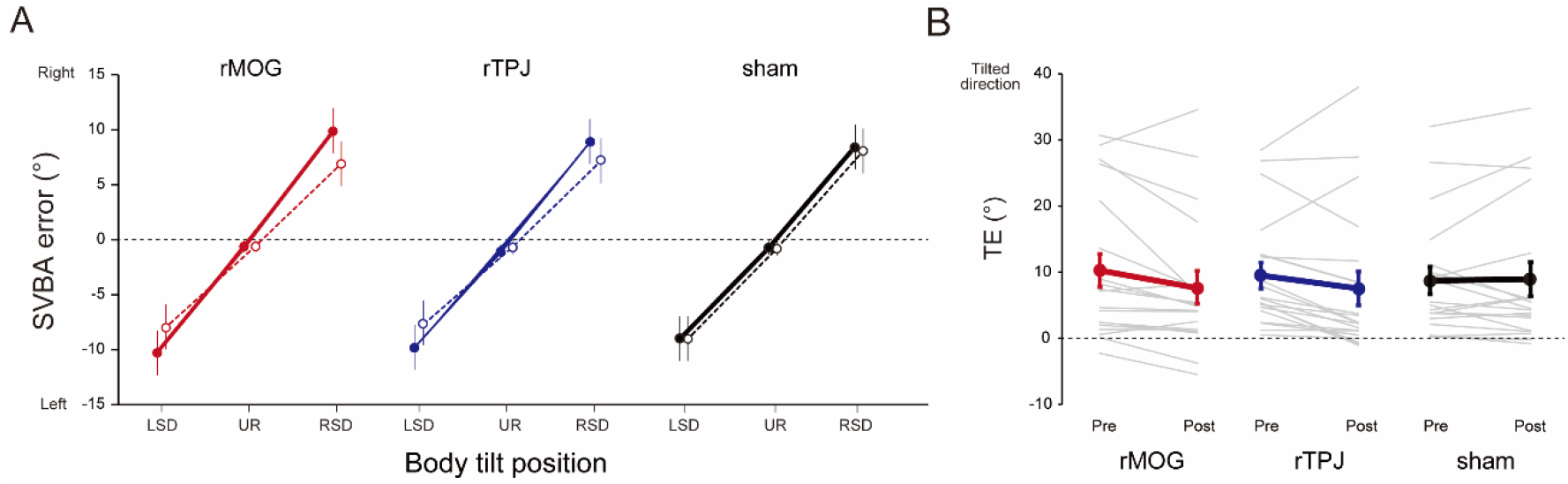
**(A)** Group-mean SVBA error for the Pre-(solid lines and filled markers) and Post-rTMS (dashed lines and open markers) phases in each rTMS condition. (B) The *TE* in each phase and rTMS condition. Grey thin and colored (red: rMOG, blue: rTPJ, black: sham) thick lines denote the *TE* for each participant and the mean across participants, respectively. Error bars represent standard errors.

As shown in Figure 2B, the *TE* tended to decrease after rTMS over the rMOG or rTPJ, while this was not the case in the sham condition. The mean *ΔTE* (Fig. 3) was negative for the rMOG (−2.65 ± 0.98°) and rTPJ (−1.99 ± 1.03°), but not for the sham (0.20 ± 0.84°) stimulation. A one-way ANOVA of *ΔTE* revealed that the main effect of rTMS condition was marginally significant (*F*_2, 36_ = 2.97, *p* = 0.06, partial *η*^2^ = 0.14), and Dunnett post-hoc test showed that the *ΔTE* in the rMOG condition was significantly smaller than that in the sham condition (*p* = 0.04, Cohen’s *d* = − 0.69). In contrast, no significant difference was observed between the rTPJ and sham conditions (*p* = 0.15, Cohen’s *d* = −0.53). We also confirmed that the *TE* in the Pre-rTMS phase (“Pre” in Fig. 2B) did not significantly differ between the rTMS conditions (rMOG, 10.02 ± 2.51°; rTPJ, 9.33 ± 1.99°; sham, 8.54 ± 2.07°; one-way ANOVA, main effect: *F*_2, 36_ = 0.65, *p* = 0.53, partial *η*^2^ = 0.04; Dunnett post-hoc test, rMOG vs. sham, *p* = 0.78, Cohen’s *d* = 0.15; rTPJ vs. sham, *p* = 0.80, Cohen’s *d* = 0.08). These results indicate that rTMS over the rMOG could reduce the perceptual distortion of egocentric spatial orientation induced by body tilt.

**Figure 3.**
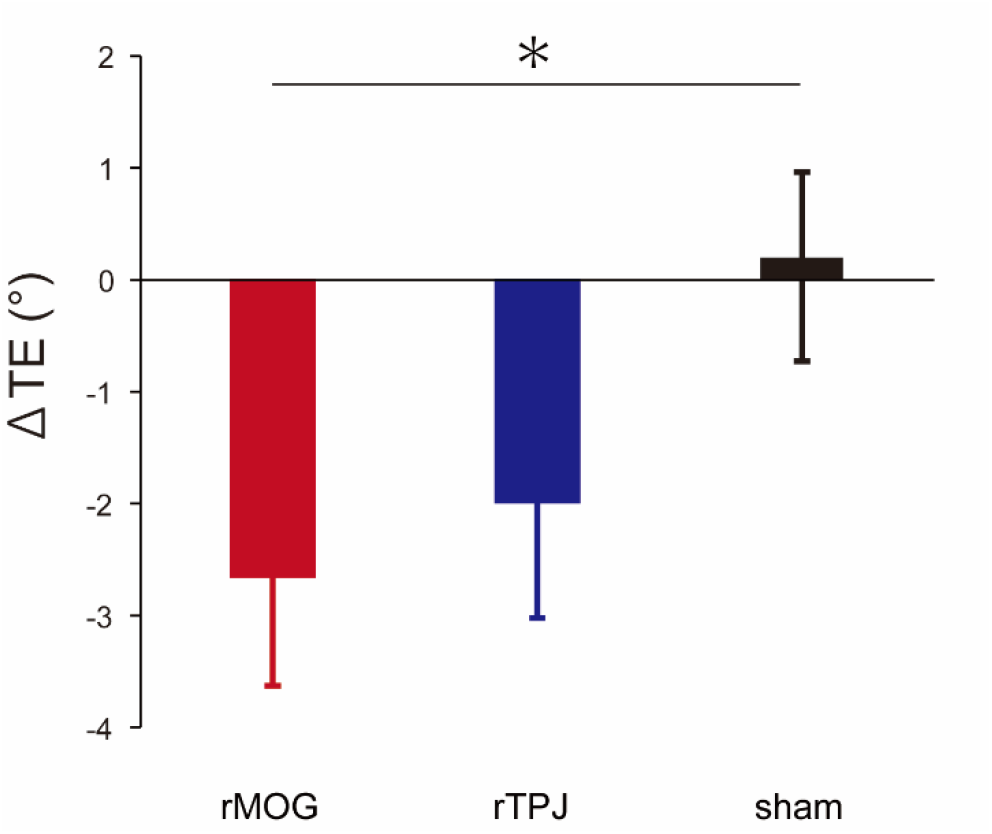
The mean *ΔTE* across participants in each rTMS condition. Error bars represent standard error. *: *p* < 0.05.

We evaluated whether the rTMS affected SVBA performance when the body was upright (*SVBAU*_*R*_), and found no significant effect of rTMS on Δ*SVBAU*_*R*_ (rMOG, −0.25 ± 0.45°; rTPJ, 0.37 ± 0.36°; sham, 0.06 ± 0.28°; ANOVA, *F*_2, 36_ = 0.65, *p* = 0.53, partial *η*^2^ = 0.04; Dunnett tests, rMOG vs. sham, *p* = 0.78, Cohen’s *d* = −0.2; rTPJ vs. sham, *p* = 0.80, Cohen’s *d* = 0.19; Fig. 4). Additionally, we examined whether the rTMS affected the trial-by-trial variability of *SVBAU*_*R*_. We calculated the standard deviation of SVBA errors across trials in the UR position (upright standard deviation: *SD*_*UR*_). As with *SVBA*_*UR*_, we calculated *ΔSD* _*UR*_ by subtracting *SD*_*UR*_ in the Pre-rTMS phase from that in the Post-rTMS phase for each rTMS condition, and performed a one-way ANOVA for *ΔSD*_*UR*_. We found that *ΔSD*_*UR*_ did not significantly differ between the rTMS conditions (rMOG, 0.04 ± 0.09°; rTPJ, −0.10 ± 0.18°; sham, 0.14 ± 0.07°; ANOVA, *F*_2, 36_ = 0.88, *p* = 0.42, partial *η*^2^ = 0.05; Dunnett tests, rMOG vs. sham, *p* = 0.62, Cohen’s *d* = −0.16; rTPJ vs. sham, *p* = 0.37, Cohen’s *d* = −0.43). These results indicate that rTMS over the rMOG did not influence SVBA performance when the body was upright.

**Figure 4.**
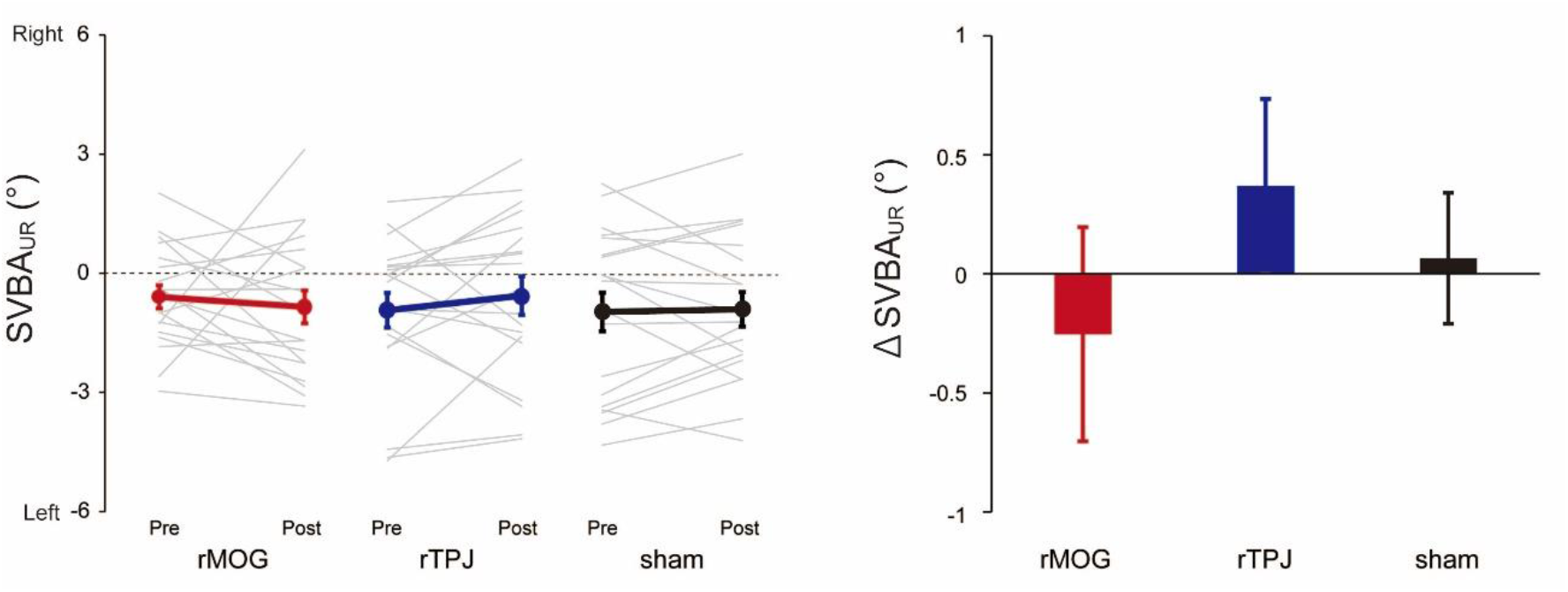
(A) The *SVBAU*_*R*_ in each phase and rTMS condition. Grey thin and colored (red: rMOG, blue: rTPJ, black: sham) thick lines denote the *SVBAU*_*R*_ for each participant and the mean across participants, respectively. Error bars represent standard error. (B) The mean Δ *SVBAU*_*R*_ across participants in each rTMS condition. Error bars represent standard error.

## Discussion

In this study, we used a whole-body tilt device and low-frequency (1 Hz) rTMS to validate the causality of neural processing in the rMOG with egocentric spatial orientation when the body is tilted. We found that perceptual distortions (measured as TE) towards the tilted side of the body in the SVBA task significantly decreased after rTMS over the rMOG. Given that 1-Hz rTMS leads to suppression of neural processing at the stimulated cortical site (Romero et al., 2002; Fitzgerald et al., 2006), our results indicate that when neural processing in the rMOG is suppressed, the body-tilt-induced distortion of egocentric spatial orientation is reduced.

Previously, we demonstrated a significant association between grey matter volume in the rMOG and TE in the SVBA task using voxel-based morphometry (Tani and Tanaka, 2021). Importantly, we observed a positive correlation: a smaller rMOG volume was associated with a smaller TE. Given that numerous studies show that greater grey matter volume is related to better functionality in a given brain region (e.g., Maguire et al., 2000), this positive correlation suggests that lower rMOG functionality leads to smaller TE. This is supported by our recent neuropsychological study in hemispheric stroke patients, which showed that lesions in the right occipitotemporal cortex, including the rMOG, were associated with an abnormally small TE (Tani et al., 2023c). In light of the two previous studies mentioned above, the significant reduction in TE observed in the current study might be attributable to the rTMS-induced suppression of neural activity in the rMOG resulting from rTMS, thereby reducing the rMOG functionality.

The MOG, as part of the lateral occipital complex, is a higher-order visual cortical region that processes spatial information about a viewed object, such as shape (Cant et al., 2009), location (Liu et al., 2017; Galati et al., 2001; Saj et al., 2014; Chen et al., 2012; Reiner et al., 2010), and orientation (Leplaideur et al., 2021; Saj et al., 2019). A recent meta-analysis (Derbie et al., 2021) showed that the rMOG is activated when a task requires visuospatial processing of information about an object in reference to the participants’ own body (egocentric spatial coding), rather than to other objects (allocentric spatial coding). In the present study, we did not observe a significant effect of rTMS over the rMOG on task performance when the body was upright (SD_UR_ and SVBA_UR_; Fig. 4), but on the body-tilt-induced perceptual distortion (TE). As described in the introduction, TE in the SVBA task is likely attributable to computational process in which the brain refers to the direction of gravity for egocentric spatial orientation (Tani et al., 2023b, 2023c). Tarnutzer and colleagues (2012) proposed that a visual object’s orientation relative to the body may be computed by combining and comparing information about the orientation of body and a visual object in relation to gravity, which is predominantly derived from otolith signals. If this is the case, the present results suggest that the rMOG primarily contributes to the integration of visual and body tilt-dependent gravitational information to internally estimate the egocentric orientation of visual objects.

In support of this, the neural visual response of the MOG has been shown to be modulated by vestibular inputs (Deutschländer et al., 2002; Della-Justina et al., 2015; Brandt et al., 2006). Deutschländer et al. (2002) reported that simultaneous vestibular stimulation suppressed the neural response of the MOG to visual stimulation, suggesting that the MOG receives and somehow integrates vestibular signals, that is a major modality for conveying gravitational information, with visual signals. Also, in non-human primates, neural activity in response to visual inputs in the lower visual cortex, including V1 (Horn et al., 1972; Tomko et al., 1981) and V2/3 (Sauvan and Peterhans, 1999), is found to be affected by the direction of gravity. Furthermore, higher association areas such as the caudal intraparietal area (Rosenberg and Angelaki, 2014) and inferotemporal cortex (Emonds et al., 2023) have been found to visually encode the tilt orientation of an object with respect to the gravitational vertical. When combining the present results with previous neurophysiological findings, it is possible to speculate that orientation information about visual objects might be integrated with vestibular-derived gravitational information at multiple stages of bottom-up visual processing, including the MOG.

As for the rTPJ, we cannot draw conclusive statements about the role of this region in integrating vestibular-derived gravitational information for egocentric spatial orientation because we observed the trend of TE reduction by rTMS to the rTPJ despite the lack of significance (Figs. 2 and 3). Further studies with larger sample sizes are needed to determine whether the observed trend reflects neural processing in the rTPJ or merely transient fluctuation.

In conclusion, our pre-registered TMS study shows, for the first time, that neural processing in the right middle occipital gyrus is an essential cause of body-tilt-induced distortion of egocentric spatial orientation, suggesting a role of the rMOG in the estimation of egocentric orientation of visual objects in reference to gravitational information. However, future behavioral and neurophysiological studies are required to fully understand the computational processes underlying the present findings.

## Acknowledgments

We are grateful to Dr. Satoshi Tanaka from Hamamatsu University School of Medicine for the technical assistance and helpful advice regarding this study. This study was supported by the Japan Society for the Promotion of Science (JSPS) KAKENHI (20K19305 and 23K16761) and Otemon Gakuin University (all to K.T.).

## References

Barra J, Oujamaa L, Chauvineau V, Rougier P, Perennou D (2009) Asymmetric standing posture after stroke is related to a biased egocentric coordinate system. Neurology 72:1582–1587.

Barra J, Benaim C, Chauvineau V, Ohlmann T, Gresty M, Perennou D (2008) Are rotations in perceived visual vertical and body axis after stroke caused by the same mechanism? Stroke 39:3099–3101.

Bauermeister M (1964) Effect of Body Tilt on Apparent Verticality, Apparent Body Position, and Their Relation. J Exp Psychol 67:142–147.

Brandt T, Glasauer S, Stephan T, Bense S, Yousry TA, Deutschlander A, Dieterich M (2002) Visual-vestibular and visuovisual cortical interaction: new insights from fMRI and pet. Ann N Y Acad Sci 956:230–241.

Cant JS, Arnott SR, Goodale MA (2009) fMR-adaptation reveals separate processing regions for the perception of form and texture in the human ventral stream. Exp Brain Res 192:391–405.

Ceyte H, Cian C, Trousselard M, Barraud PA (2009) Influence of perceived egocentric coordinates on the subjective visual vertical. Neurosci Lett 462:85–88.

Ceyte H, Cian C, Nougier V, Olivier I, Trousselard M (2007) Role of gravity-based information on the orientation and localization of the perceived body midline. Experimental Brain Research 176:504–509.

Chen Q, Weidner R, Weiss PH, Marshall JC, Fink GR (2012) Neural interaction between spatial domain and spatial reference frame in parietal-occipital junction. J Cogn Neurosci 24:2223–2236.

Clement G, Arnesen TN, Olsen MH, Sylvestre B (2007) Perception of longitudinal body axis in microgravity during parabolic flight. Neurosci Lett 413:150–153.

Clifford CW, Webster MA, Stanley GB, Stocker AA, Kohn A, Sharpee TO, Schwartz O (2007) Visual adaptation: neural, psychological and computational aspects. Vision Res 47:3125–3131.

Della-Justina HM, Gamba HR, Lukasova K, Nucci-da-Silva MP, Winkler AM, Amaro E, Jr. (2015) Interaction of brain areas of visual and vestibular simultaneous activity with fMRI. Exp Brain Res 233:237–252.

Derbie AY, Chau BKH, Wong CHY, Chen LD, Ting KH, Lam BYH, Lee TMC, Chan CCH, Smith Y (2021) Common and distinct neural trends of allocentric and egocentric spatial coding: An ALE meta-analysis. Eur J Neurosci 53:3672–3687.

Deutschlander A, Bense S, Stephan T, Schwaiger M, Brandt T, Dieterich M (2002) Sensory system interactions during simultaneous vestibular and visual stimulation in PET. Hum Brain Mapp 16:92–103.

Donaldson PH, Rinehart NJ, Enticott PG (2015) Noninvasive stimulation of the temporoparietal junction: A systematic review. Neurosci Biobehav Rev 55:547–572.

Emonds AMX, Srinath R, Nielsen KJ, Connor CE (2023) Object representation in a gravitational reference frame. Elife 12.

Fiori F, Candidi M, Acciarino A, David N, Aglioti SM (2015) The right temporoparietal junction plays a causal role in maintaining the internal representation of verticality. J Neurophysiol 114:2983–2990.

Fitzgerald PB, Fountain S, Daskalakis ZJ (2006) A comprehensive review of the effects of rTMS on motor cortical excitability and inhibition. Clin Neurophysiol 117:2584–2596.

Galati G, Committeri G, Sanes JN, Pizzamiglio L (2001) Spatial coding of visual and somatic sensory information in body-centred coordinates. Eur J Neurosci 14:737–746.

Horn G, Hill RM (1969) Modifications of Receptive Fields of Cells in the Visual Cortex occurring Spontaneously and associated with Bodily Tilt. Nature 221:186–188.

Hothorn LA (2016) The two-step approach—a significant ANOVA F-test before Dunnett’s comparisons against a control—is not recommended. Communications in Statistics - Theory and Methods 45:3332–3343.

Jamal K, Leplaideur S, Rousseau C, Chochina L, Moulinet-Raillon A, Bonan I (2018) Disturbances of spatial reference frame and postural asymmetry after a chronic stroke. Exp Brain Res 236:2377–2385.

Kassuba T, Klinge C, Holig C, Roder B, Siebner HR (2014) Short-term plasticity of visuo-haptic object recognition. Front Psychol 5:274.

Kheradmand A, Lasker A, Zee DS (2015) Transcranial magnetic stimulation (TMS) of the supramarginal gyrus: a window to perception of upright. Cereb Cortex 25:765–771.

Leplaideur S, Moulinet-Raillon A, Duche Q, Chochina L, Jamal K, Ferre JC, Bannier E, Bonan I (2021) The Neural Bases of Egocentric Spatial Representation for Extracorporeal and Corporeal Tasks: An fMRI Study. Brain Sci 11.

Liu N, Li H, Su W, Chen Q (2017) Common and specific neural correlates underlying the spatial congruency effect induced by the egocentric and allocentric reference frame. Hum Brain Mapp 38:2112–2127.

Maguire EA, Gadian DG, Johnsrude IS, Good CD, Ashburner J, Frackowiak RS, Frith CD (2000) Navigation-related structural change in the hippocampi of taxi drivers. Proc Natl Acad Sci U S A 97:4398–4403.

Mars F, Vercher JL, Popov K (2005) Dissociation between subjective vertical and subjective body orientation elicited by galvanic vestibular stimulation. Brain Res Bull 65:77–86.

McFarland JH, Clarkson F (1966) PERCEPTION OF ORIENTATION - ADAPTATION TO LATERAL BODY-TILT. American Journal of Psychology 79:265-+.

Mills KR, Boniface SJ, Schubert M (1992) Magnetic brain stimulation with a double coil: the importance of coil orientation. Electroencephalogr Clin Neurophysiol 85:17–21.

Nicholls ME, Thomas NA, Loetscher T, Grimshaw GM (2013) The Flinders Handedness survey (FLANDERS): a brief measure of skilled hand preference. Cortex 49:2914–2926.

Nosek BA, Ebersole CR, DeHaven AC, Mellor DT (2018) The preregistration revolution. Proc Natl Acad Sci U S A 115:2600–2606.

Renier LA, Anurova I, De Volder AG, Carlson S, VanMeter J, Rauschecker JP (2010) Preserved functional specialization for spatial processing in the middle occipital gyrus of the early blind. Neuron 68:138–148.

Romero JR, Anschel D, Sparing R, Gangitano M, Pascual-Leone A (2002) Subthreshold low frequency repetitive transcranial magnetic stimulation selectively decreases facilitation in the motor cortex. Clin Neurophysiol 113:101–107.

Rosenberg A, Angelaki DE (2014) Gravity influences the visual representation of object tilt in parietal cortex. J Neurosci 34:14170–14180.

Saj A, Borel L, Honore J (2019) Functional Neuroanatomy of Vertical Visual Perception in Humans. Front Neurol 10:142.

Saj A, Cojan Y, Musel B, Honore J, Borel L, Vuilleumier P (2014) Functional neuro-anatomy of egocentric versus allocentric space representation. Neurophysiol Clin 44:33–40.

Santos-Pontelli TE, Rimoli BP, Favoretto DB, Mazin SC, Truong DQ, Leite JP, Pontes-Neto OM, Babyar SR, Reding M, Bikson M, Edwards DJ (2016) Polarity-Dependent Misperception of Subjective Visual Vertical during and after Transcranial Direct Current Stimulation (tDCS). PLoS One 11:e0152331.

Sauvan XM, Peterhans E (1999) Orientation constancy in neurons of monkey visual cortex. Visual Cognition 6:43–54.

Tamura A, Wada Y, Inui T, Shiotani A (2017) Perceived direction of gravity and the body-axis during static whole body roll-tilt in healthy subjects. Acta Otolaryngol 137:1057–1062.

Tani K, Tanaka S (2021) Neuroanatomical correlates of the perception of body axis orientation during body tilt: a voxel-based morphometry study. Sci Rep 11:14659.

Tani K, Uehara S, Tanaka S (2023a) Psychophysical evidence for the involvement of head/body-centered reference frames in egocentric visuospatial memory: A whole-body roll tilt paradigm. J Vis 23:16.

Tani K, Uehara S, Tanaka S (2023b) Association Between Body Tilt and Egocentric Estimates Near Upright. Multisens Res 36:367–386.

Tani K, Shiraki Y, Yamamoto S, Kodaka Y, Kushiro K (2018) Whole-body roll tilt influences goal-directed upper limb movements through the perceptual tilt of egocentric reference frame. Frontiers in Psychology 9:84.

Tani K, Iio S, Kamiya M, Yoshizawa K, Shigematsu T, Fujishima I, Tanaka S (2023c) Neuroanatomy of reduced distortion of body-centred spatial coding during body tilt in stroke patients. Sci Rep 13:11853.

Tarnutzer AA, Bockisch CJ, Olasagasti I, Straumann D (2012) Egocentric and allocentric alignment tasks are affected by otolith input. Journal of Neurophysiology 107:3095–3106.

Tomko DL, Barbaro NM, Ali FN (1981) Effect of body tilt on receptive field orientation of simple visual cortical neurons in unanesthetized cats. Exp Brain Res 43:309–314.

Tsakiris M, Costantini M, Haggard P (2008) The role of the right temporo-parietal junction in maintaining a coherent sense of one’s body. Neuropsychologia 46:3014–3018.

